# High-throughput, label-free and slide-free histological imaging by computational microscopy and unsupervised learning

**DOI:** 10.1101/2021.06.04.447030

**Authors:** Yan Zhang, Lei Kang, Xiufeng Li, Ivy H. M. Wong, Terence T. W. Wong

## Abstract

Rapid and high-resolution histological imaging with minimal tissue preparation has long been a challenging and yet captivating medical pursue. Here, we propose a promising and transformative histological imaging method, termed computational high-throughput autofluorescence microscopy by pattern illumination (CHAMP). With the assistance of computational microscopy, CHAMP enables high-throughput and label-free imaging of thick and unprocessed tissues with large surface irregularity at an acquisition speed of 10 mm^2^/10 seconds with 1.1-µm lateral resolution. Moreover, the CHAMP image can be transformed into a virtually stained histological image (Deep-CHAMP) through unsupervised learning within 15 seconds, where significant cellular features are quantitatively extracted with high accuracy. The versatility of CHAMP is experimentally demonstrated using mouse brain/kidney tissues prepared with various clinical protocols, which enables a rapid and accurate intraoperative/postoperative pathological examination without tissue processing or staining, demonstrating its great potential as an assistive imaging platform for surgeons and pathologists to provide optimal adjuvant treatment.

## Introduction

Postoperative histological examination by pathologists remains the gold standard for surgical margin assessment (SMA), which aims to examine if there are any remaining cancer cells at the cut margin^1^. However, routine pathological examination based on formalin-fixed and paraffin-embedded (FFPE) tissues involves a series of lengthy and laborious steps (Supplementary Fig. 1a), causing a significant delay (ranging from hours to days) in providing accurate diagnostic reports. Although intraoperative frozen section can serve as a rapid alternative for SMA, it suffers from freezing artifacts when dealing with edematous and soft tissues, and sub-optimal cutting for fatty tissues, affecting slide interpretation and diagnostic accuracy^2^.

**Figure 1.**
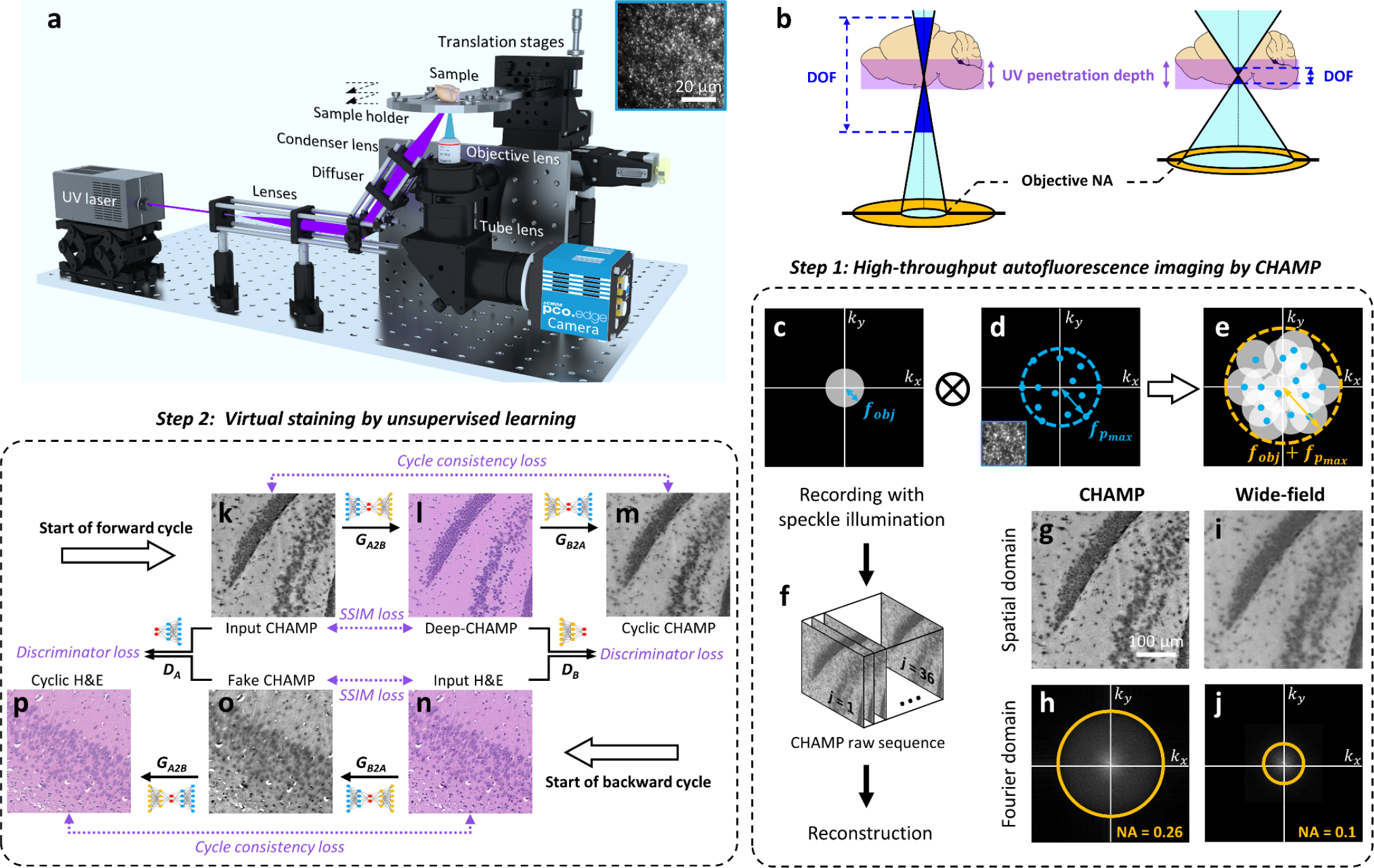
Overview of histological imaging by CHAMP. **a**, Schematic of the CHAMP system. The beam is expanded by a pair of lenses, and obliquely reflected onto a diffuser to generate an interference-induced speckle pattern (inset at the top right corner), which is subsequently focused onto the bottom surface of a specimen by a condenser lens. The excited autofluorescence signal is collected by an objective lens, refocused by a tube lens, and subsequently detected by a monochromatic camera. The specimen supported by a sample holder is raster-scanned by a 2-axis motorized stage to generate a sequence of speckle-illuminated diffraction-limited autofluorescence images. **b**, Illustration of the relationship between DOF, objective NA, UV penetration depth (i.e., UV optical-sectioning thickness), and tissue surface irregularity. **c**, The aperture of the diffraction-limited imaging system with a cut-off frequency of *f*_obj_. **d**, The spectrum of the speckle pattern with a maximum frequency of 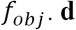 . **e**, The synthetic aperture 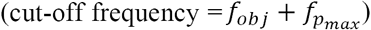 through intensity modulation by speckle illumination. **f**, The captured raw image sequence for CHAMP reconstruction. **g**,**h**, A reconstructed resolution-enhanced image (i.e., a CHAMP image) and its corresponding spectrum in Fourier domain, respectively. **i**,**j**, Diffraction-limited wide-field image captured with uniform illumination and its corresponding spectrum in the Fourier domain, respectively. **k**, Input CHAMP image. **l**, Virtually stained Deep-CHAMP image. **m**, Generated cyclic CHAMP image. **n**, Input H&E-stained image. **o**, Generated fake CHAMP image. **p**, Generated cyclic H&E-stained image.

The great demand in histopathology has inspired lots of efforts in achieving a rapid and non-invasive diagnosis for unprocessed tissues. Some cutting-edge microscopy techniques (Supplementary Fig. 2) with optical sectioning capability enable slide-free imaging of thick resection specimens, greatly simplifying the procedures associated with tissue sectioning in conventional FFPE. The scanning-based depth-resolved approaches, including confocal microscopy^3,4^, photoacoustic microscopy (PAM)^5,6^, multiphoton microscopy (MPM)^7^, stimulated Raman scattering (SRS)^8,9^, second harmonic generation (SHG)^10^, and their spectral multiplexing^11,12^, enables surface profiling of bulk tissues via two-dimensional/three-dimensional (2D)/(3D) scanning of a tightly focused laser beam. However, the imaging throughput is therefore restricted to tens of megapixels (Supplementary Fig. 2) due to the low repetition rate of pulsed lasers and long pixel dwell time, posing a challenge to examine large specimens (e.g., human biopsies) with a centimeter-scale surface area within a short diagnostic time frame. In contrast, wide-field depth-resolved techniques, including microscopy with ultraviolet surface excitation (MUSE)^13,14^, light-sheet microscopy^15,16^, and structured illumination microscopy (SIM)^17–20^, are fundamentally suitable for time-sensitive applications as their imaging throughput can reach hundreds of megapixels via parallel pixel acquisition. However, fluorescence labelling is generally required by these methods to visualize features that analogous to FFPE histology, which is challenging to be integrated into the current clinical practice. Given this, label-free imaging contrast is highly desirable in modern clinical settings.

**Figure 2.**
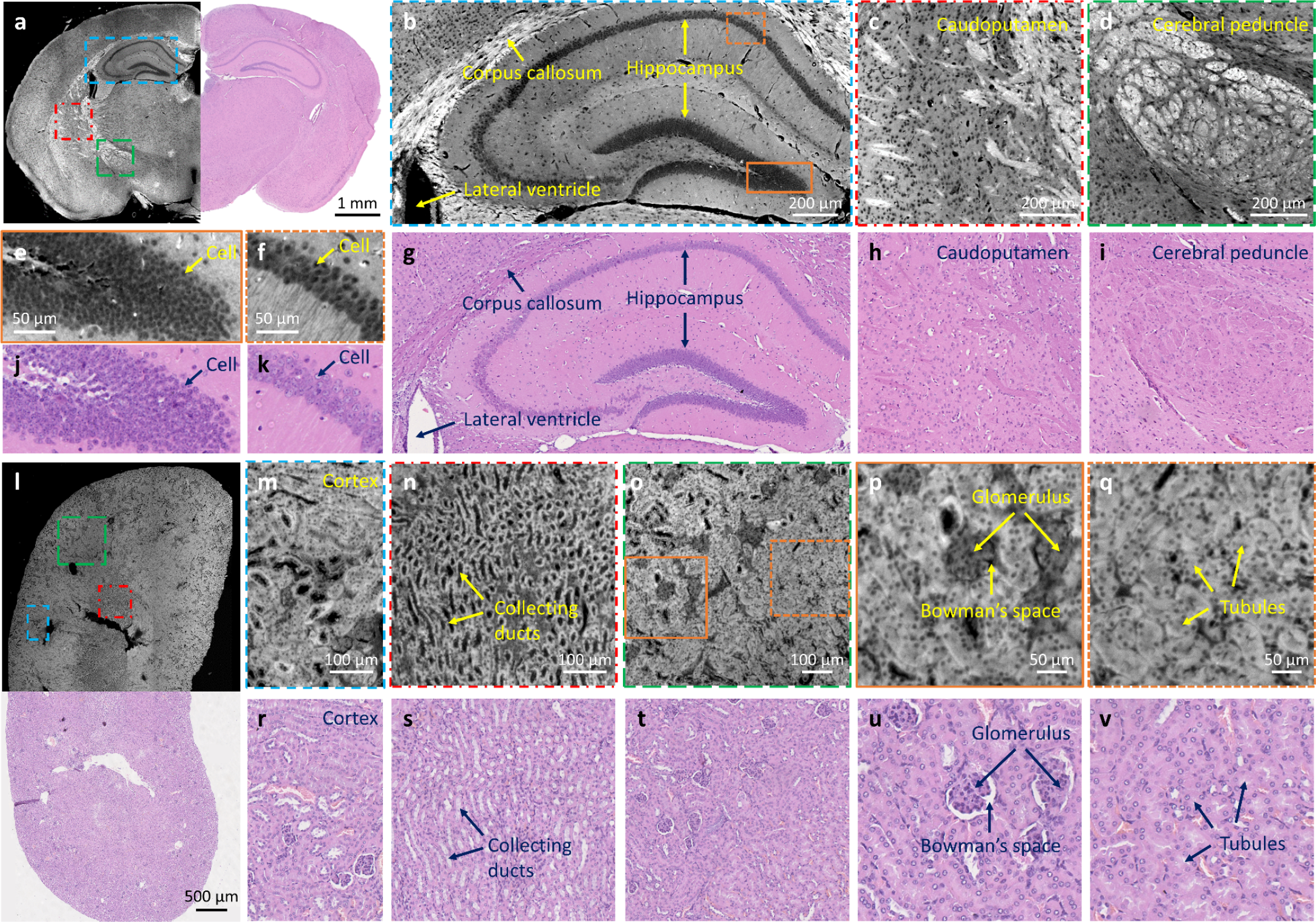
CHAMP and histological imaging of thin mouse brain/kidney tissue slices. **a**, Combined CHAMP and H&E-stained mosaic image of a mouse brain. **b**–**d**, Zoomed-in CHAMP images of blue, red, and green dashed regions in a, respectively. **e**,**f**, Zoomed-in CHAMP images of orange solid and dashed regions in b, respectively. **g**–**k**, The corresponding H&E-stained images. **l**, Combined CHAMP and H&E-stained mosaic image of a mouse kidney. **m**–**o**, Zoomed-in CHAMP images of blue, red, and green dashed regions in l, respectively. **p**,**q**, Zoomed-in CHAMP images of orange solid and dashed regions in o, respectively. **r**–**v**, The corresponding H&E-stained images.

The aforementioned PAM and nonlinear microscopy play an indispensable role in non-invasive label-free characterization of various biological structures, through either absorption-induced thermoelastic expansion (PAM), intrinsic autofluorescence (MPM), molecular vibration (SRS), or non-centrosymmetric orientation (SHG). The simultaneous multiplexing^21,22^ of these methods enables cell phenotyping and classification based on the targeted biomolecules. However, the low-throughput nature of these scanning-based imaging modalities ultimately hinders their clinical translations. In addition, reflectance-based imaging approaches, such as optical coherence tomography (OCT)^23,24^ and reflectance confocal microscopy (RCM)^25^, pave a way for rapid label-free inspection of human breast tissue. However, they are typically not designed at subcellular resolution.

Standard histological images are acquired at subcellular resolution which is essential for pathological analysis. However, the inherent trade-offs between resolution, field-of-view (FOV), and depth-of-field (DOF) fundamentally pose an impediment for rapid and high-resolution imaging of thick tissues through a high numerical aperture (NA) objective lens. Firstly, the image quality will be significantly degraded once the resulting shallow DOF (typically a few microns) is shorter than the optical-sectioning thickness of the employed imaging modality, which is tunable with light-sheet microscopy and SIM, but tissue-dependent with MUSE (determined by the ultraviolet (UV) penetration depth (∼100 µm in human breast^5^ and ∼20 µm in human skin^26^)). In addition, the shallow DOF is unable to accommodate various surface irregularities and tissue debris presented in surgical resection specimens, leading to severe out-of-focus blurs which ultimately prevent high-quality imaging of fine structures in thick specimens. Although extended DOF^27^ can be applied to extract in-focus information at the tissue surface through a sequence of axially-refocused images, the achievable throughput of the system is largely sacrificed.

Here, we propose a promising and transformative histological imaging technology, termed computational high-throughput autofluorescence microscopy by pattern illumination (CHAMP), which enables high-throughput and label-free imaging of thick and unprocessed tissues with large surface irregularity at an acquisition speed of 10 mm^2^/10 seconds with 1.1-µm lateral resolution. To the best of our knowledge, this is not achievable with any of the existing methods. Rich endogenous fluorophores^28^, including reduced nicotinamide adenine dinucleotide, structural proteins (e.g., collagen and elastin), aromatic amino acids (e.g., tryptophan, tyrosine), and lipofuscin, naturally form a fundamental contrast mechanism with deep-UV excitation in CHAMP. High imaging throughput and long DOF can be achieved with the assistance of computational microscopy, making CHAMP highly suitable for intraoperative tissue assessment (e.g., SMA) where immediate feedback should be provided to surgeons for optimal adjuvant treatment. Furthermore, an unsupervised neural network is implemented to transform a CHAMP image of an unlabeled tissue into a virtually stained histological image (Deep-CHAMP), ensuring an easy interpretation by pathologists. As thick resection tissues are inevitably deformed during the FFPE workflow (e.g., rigidity change, tissue shrinkage, tissue rupture, and slide folding), it is impractical to obtain a pixel-to-pixel registered label-free CHAMP image with its corresponding histological image to form a paired training data as required by some recently reported virtual staining networks^29,30^. In contrast, we employ unsupervised learning based on the architecture of a cycle-consistent generative adversarial network (CycleGAN)^31^, which enables image translation without paired training data, fundamentally favouring the virtual staining of CHAMP images of thick and unprocessed tissues. Diagnostic features are quantitatively extracted from Deep-CHAMP with high accuracy. The versatility of CHAMP is experimentally demonstrated using mouse brain/kidney tissues prepared with various clinical protocols, which enables rapid and accurate intraoperative/postoperative pathological examinations without tissue processing or staining. The high-throughput, high-versatility, cost-effective, and ease-of-use features of our CHAMP microscope holds great promise in clinical translations to revolutionize the current gold standard histopathology.

## Results

### Histological imaging by CHAMP microscopy

Our CHAMP microscope is configured in a reflection mode (Fig. 1a and Supplementary Video 1), which can accommodate tissues with any size and thickness without physically interfering with the illumination and collection optics. Deep-UV laser at 266 nm, which presents significant difference of the quantum yields between nucleotides^32^ and other endogenous fluorophores (e.g., tryptophan), is used for illumination in our CHAMP system to maximize the negative contrast of cell nuclei. Oblique illumination circumvents the use of UV-transmitting optics and fluorescence filters because the backscattered UV light is naturally blocked by the glass objective and tube lens which are spectrally opaque at 266 nm. A constant speckle pattern (inset of Fig. 1a), which is generated by a diffuser and featured a grain size smaller than the point spread function (PSF) of the detection optics, is projected onto the bottom surface of the specimen for pattern illumination. A long DOF (Fig. 1b) enabled by the implementation of a low-NA objective lens not only matches the optical-sectioning thickness provided by UV surface excitation, but also accommodates different levels of tissue surface irregularities. With intensity modulation, the aperture of the diffraction-limited system (Fig. 1c) is convolved with the spectrum of speckle pattern which contains various frequency components (Fig. 1d), and is consequently 2D translated in the Fourier domain, enabling the synthesis of an extended system passband (Fig. 1e). This allows high spatial frequency (i.e., high-resolution features) to be encoded into the low-NA imaging system, thus bypassing the resolution limit governed by the low-NA objective lens equipped in the CHAMP microscope. The sample is raster-scanned to generate a sequence of speckle-illuminated diffraction-limited images (Fig. 1f), which are subsequently demodulated to reconstruct a resolution-enhanced image (Fig. 1g and Supplementary Video 1) (termed CHAMP image hereafter). CHAMP imaging features 2.6× resolution improvement (see Methods) compared with conventional wide-field microscopy with uniform illumination (Fig. 1i). The reconstructed CHAMP image is subsequently transformed into a virtually stained histological image (termed Deep-CHAMP image hereafter) through a CycleGAN-based neural network, which is composed of four deep neural networks, including two generators (*G*_*A2B*_, *G*_*B2A*_) and two discriminators (*D*_*A*_, *D*_*B*_). The generator *G*_*A2B*_ learns to transform grayscale images to color images, while the generator *G*_*B2A*_ learns to transform color images to grayscale images. A sequence of unpaired CHAMP images and hematoxylin and eosin (H&E) stained images, are fed to the neural network to undergo a forward training cycle (Fig. 1k–m) and a backward training cycle (Fig. 1n–p). The discriminator *D*_*A*_ aims to distinguish real input CHAMP images (Fig. 1k) from fake CHAMP images (Fig. 1o) produced by the generator *G*_*B2A*_. Meanwhile, the discriminator *D*_*B*_ aims to distinguish real input H&E-stained images (Fig. 1n) from virtually stained Deep-CHAMP images (Fig. 1l) produced by the generator *G*_*A2B*_. Once the generator *G*_*A2B*_ can produce Deep-CHAMP images that the discriminator *D*_*B*_ cannot distinguish them from the input H&E-stained images, the transformation from CHAMP to Deep-CAHMP is well learned by the generator *G*_*A2B*_. This iterative process is also applicable to the generator *G*_*B2A*_ and the discriminator *D*_*A*_.

### CHAMP and histological imaging of thin mouse brain/kidney tissue slices

FFPE thin slices of mouse brain/kidney tissues are imaged to validate the performance of CHAMP initially (Fig. 2). The microtome-sectioned thin slices (with thickness ∼7 µm) are deparaffinized before CHAMP imaging. Cell nuclei distributed at cerebral cortex and brain stem are clearly revealed with a negative contrast in CHAMP images. With a measured resolution of 1.1 µm (see Methods), the densely packed cell nuclei in the hippocampus (Fig. 2b, and zoomed-in CHAMP images (Fig. 2e,f) of orange solid and dashed regions in Fig. 2b) can be resolved individually. Other anatomical structures, including lateral ventricle and corpus callosum (Fig. 2b), caudoputamen (Fig. 2c), and cerebral peduncle (Fig. 2d) are also well recognized. After CHAMP imaging, the slice is histologically stained by H&E, and imaged with a bright-field microscope to obtain the corresponding histological images (Fig. 2g–k). The cerebral peduncle, which is poorly visualized in the H&E-stained image (Fig. 2i), can be clearly identified in CHAMP (Fig. 2d). Multiple similarities are revealed in CHAMP and H&E-stained images, despite that the nucleoli are less visible in CHAMP. Pearson correlation coefficient of 0.9 is calculated from Fig. 2b and Fig. 2g, validating the feasibility of using tissue’s autofluorescence as an intrinsic contrast mechanism for label-free characterization of biological structures. Similarly, CHAMP provides well-characterized structures of a mouse kidney (Fig. 2l–q), including cortex (Fig. 2m), collecting ducts (Fig. 2n), glomerulus and Bowman’s space (Fig. 2p), and renal tubules (Fig. 2q). Their corresponding H&E-stained images are shown in Fig. 2r–v. A reduced correlation coefficient of 0.7 is calculated from Fig. 2p and Fig. 2u, which is due to the locally deformed Bowman’s space during the subsequent H&E staining after CHAMP imaging.

### CHAMP imaging of thick and unprocessed mouse brain/kidney tissues

To showcase the slide-free and label-free imaging capability of CHAMP as well as its superiority in thick tissue imaging, formalin-fixed and unprocessed thick mouse brain/kidney tissues are imaged (Fig. 3). The mouse brain tissues are hand-cut at different coronal planes with thickness ∼5 mm, while the mouse kidney tissue is vibratome-sectioned with thickness ∼200 µm. As mentioned above, resolution-enhanced and all-in-focus (due to the long DOF) CHAMP images (Fig. 3a–d) eliminate any image blur that potentially originated from mismatched DOF and UV penetration depth or non-flattened tissue surface. Zoomed-in CHAMP images of cell nuclei in the hippocampus (Fig. 3e,g) and lobules (Fig. 3i) far outperform the corresponding wide-field images which are directly captured with a 0.3-NA imaging objective (Fig. 3f,h,j). The out-of-focus blurs presented in kidney tubules (Fig. 3m,n) are eliminated in CHAMP images (Fig. 3k,l) such that individual cells are clearly observed in the entire FOV.

**Figure 3.**
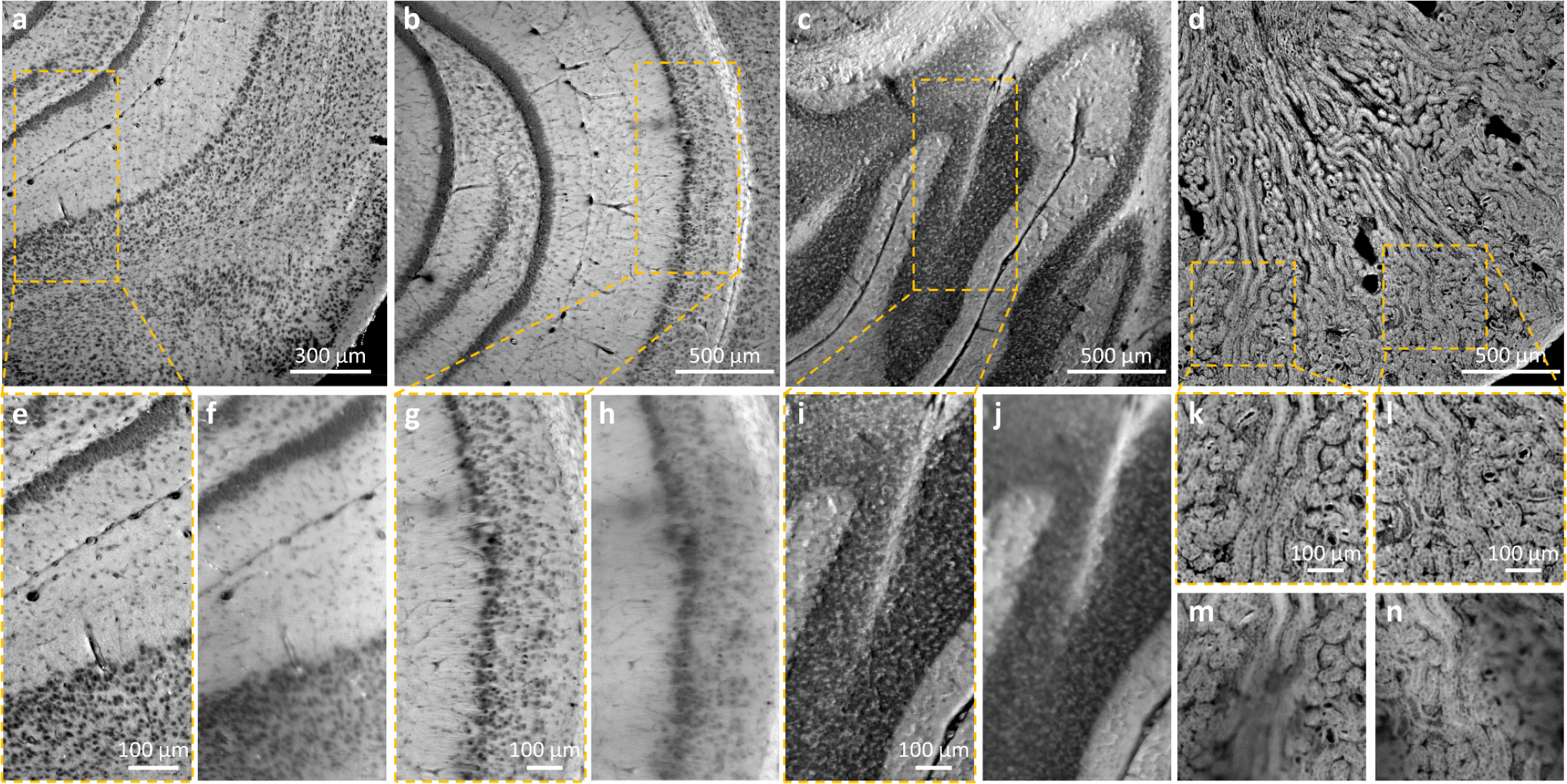
CHAMP imaging of thick and unprocessed mouse brain/kidney tissues. **a**–**c**, CHAMP images of fixed and unprocessed mouse brain tissues hand-cut at different coronal planes with thickness ∼5 mm. **d**, CHAMP image of a fixed and unprocessed mouse kidney tissue sectioned with thickness ∼200 µm. **e**,**g**,**i**,**k**,**l**, Zoomed-in CHAMP images of orange dashed regions in a**–**d. **f**,**h**,**j**,**m**,**n**, The corresponding wide-field images captured with a 0.3-NA imaging objective with uniform illumination.

### CHAMP and Deep-CHAMP imaging of thick tissues treated with various clinical protocols

CHAMP and Deep-CHAMP imaging is experimentally validated with mouse brain/kidney tissues which are treated with various clinical protocols, e.g., microtome-sectioned thin tissue slice (Supplementary Fig. 3 and Supplementary Video 2), formalin-fixed thick tissues (Fig. 4 and Supplementary Video 3, Supplementary Fig. 4 and Supplementary Video 4), as well as freshly excised tissues (Fig. 5, Supplementary Fig. 5). We trained two neural networks to separately handle the virtual staining of fixed and fresh mouse brains due to the significant difference in CHAMP images. In addition, we found the overall trend for the CycleGAN is that it converts brighter regions to white background, and darker regions to purple nuclei. Therefore, dark features in CHAMP (e.g., interstitial spaces, ventricles, and vessels) can be incorrectly color mapped to purple and mixed with cells. To alleviate this issue, the CHAMP image is segmented by a pre-trained classifier to separate cell nuclei from features that demonstrate similar brightness. After that, the segmented CHAMP image is cropped and fed into the network to output a virtually stained Deep-CHAMP image.

**Figure 4.**
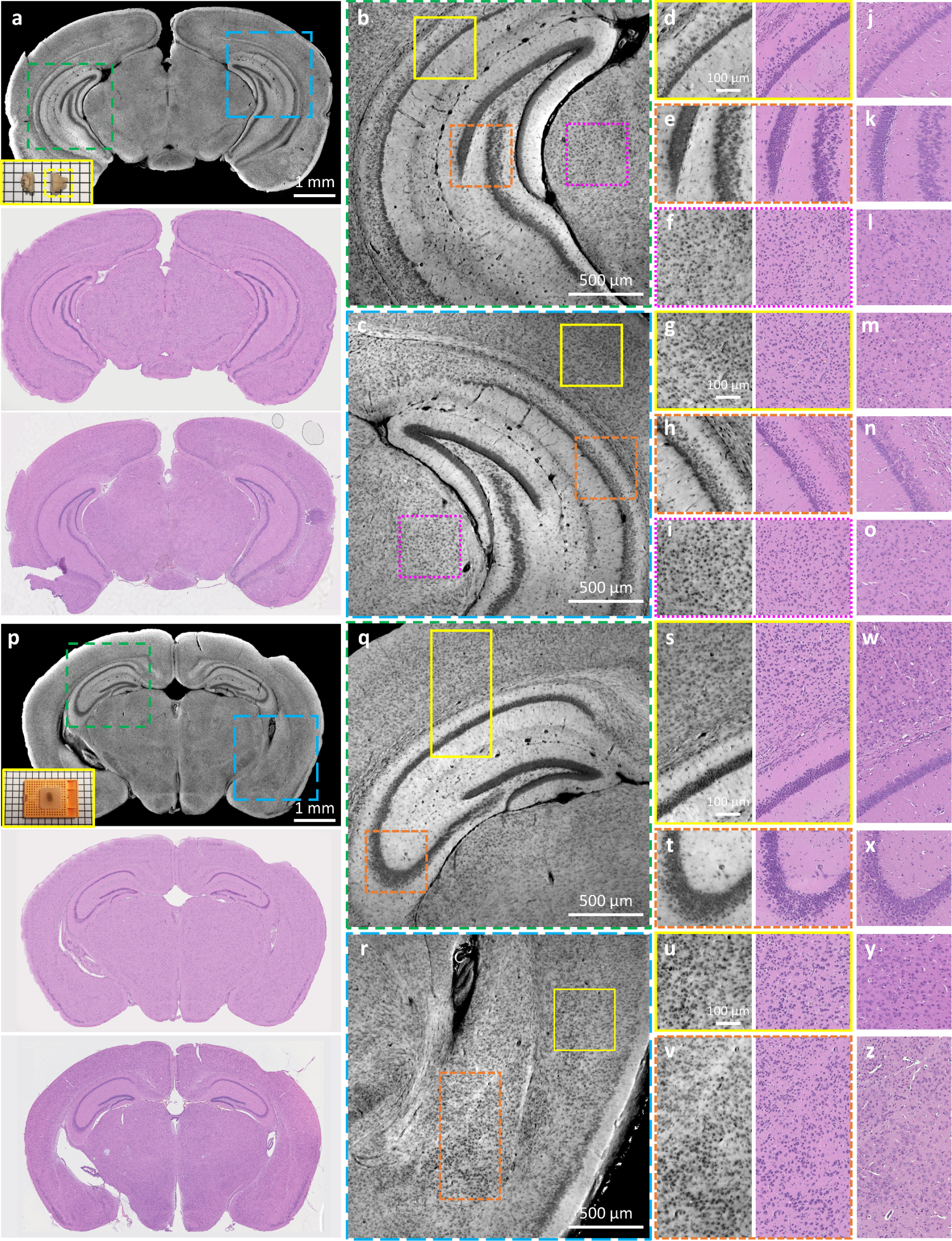
CHAMP and Deep-CHAMP validation with formalin-fixed thick mouse brains. **a**, CHAMP (top), Deep-CHAMP (middle), and H&E-stained image (bottom) of a fixed and unprocessed mouse brain, inset at the bottom left of CHAMP shows the photograph of the specimen (the yellow dashed box shows the mouse brain that is imaged). **b**,**c**, Zoomed-in CHAMP images of green and blue dashed regions in a, respectively. **d**–**f**,**g**–**i**, Zoomed-in CHAMP and Deep-CAHMP images of yellow solid, orange dashed, and magenta dashed regions in b and c, respectively. **j**–**o**, The corresponding H&E-stained images. **p**, CHAMP (top), Deep-CHAMP (middle) and H&E-stained image (bottom) of a fixed and paraffin-embedded mouse brain, inset at the bottom left of CHAMP shows the photograph of the specimen. **q**,**r**, Zoomed-in CHAMP images of green and blue dashed regions in p, respectively. **s**–**v**, Zoomed-in CHAMP and Deep-CHAMP images of yellow solid and orange dashed regions in q and r, respectively. **w**–**z**, The corresponding H&E-stained images.

**Figure 5.**
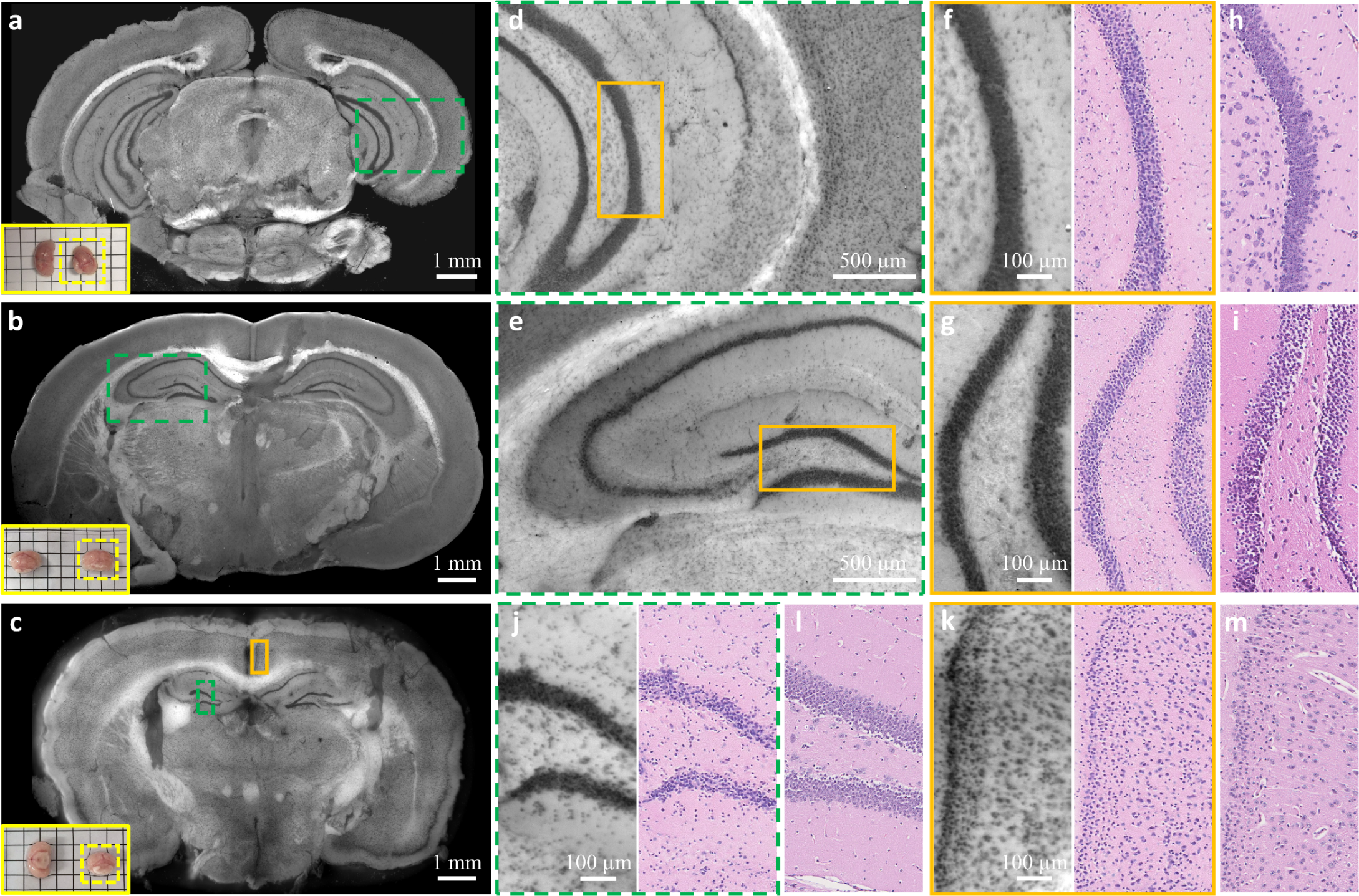
CHAMP and Deep-CHAMP validation with freshly excised mouse brains. **a**–**c**, CHAMP images of freshly excised mouse brains, inset at the bottom left of each figure shows the photograph of the specimen (the yellow dashed box indicates the mouse brain that is imaged). **d**,**e**, Zoomed-in CHAMP images of green dashed regions in a and b, respectively. **f**,**g**, Zoomed-in CHAMP and Deep-CHAMP images of orange regions in d and e, respectively. **h**,**i**, The corresponding H&E-stained images. **j**,**k**, Zoomed-in CHAMP and Deep-CHAMP images of green dashed and orange solid regions in c, respectively. **l**,**m**, The corresponding H&E-stained images.

In Fig. 4, the freshly excised mouse brains are fixed in formalin for 24 hours to prevent tissue degradation, after which the specimens are manually sectioned without any further processing (Fig. 4a), or processed and paraffin-embedded as a block tissue (Fig. 4p). The specimens are imaged by CHAMP and virtually stained to generate the corresponding Deep-CHAMP images (Fig. 4d–i,s–v), and are subsequently processed by a standard histological procedure to obtain the H&E-stained images for comparison (Fig. 4j– o,w–z). Note that the microtome-sectioned FFPE thin slice is not able to exactly replicate the surface imaged by CHAMP due to the tissue deformation and the difference in imaging thickness. Despite this difference, the structural features are still remarkably similar.

In Fig. 5, freshly excised mouse brains are rinsed in phosphate-buffered saline to remove adhesive blood on the cut surface, and blotted up before imaging. The specimens are imaged by CHAMP and virtually stained to generate the corresponding Deep-CHAMP images (Fig. 5f,g,j,k). After that, the specimens are processed following the standard procedure to obtain the H&E-stained images (Fig. 5h,i,l,m). Despite the tissue deformation and shrinkage of fresh brains (percentage of shrinkage is ∼40% in our experiments), the histological features are still considerably similar. It should be noted that cell nuclei located at heavily myelinated regions are partially obscured due to the strong scattering in fresh brain tissues.

In addition to mouse brain tissues, CHAMP and Deep-CHAMP imaging is also applied to fixed or fresh mouse kidneys (Supplementary Figs. 4,5). In fresh mouse kidney, the densely packed cell nuclei along kidney tubules are well identified in CHAMP (Supplementary Fig. 5b–e). These CHAMP images are firstly fed to the virtual staining network trained for fresh mouse brain to obtain ‘brain-style’ Deep-CHAMP images (Supplementary Fig. 5f–i), which are subsequently input to another unsupervised network trained for style transformation (see Methods), to generate ‘kidney-style’ Deep-CHAMP images (Supplementary Fig. 4j–m). This bridge network allows style transformation among different types of tissues without the need for retraining on specific tissue, demonstrating great simplicity and flexibility of the unsupervised neural networks. Note that intricate renal tubules and blood vessels in fresh kidney pose a challenge for feature segmentation, which consequently leads to some staining artifacts in the generated Deep-CHAMP images (indicated by the arrows in Supplementary Fig. 5).

Diagnostic features, such as cross-sectional area and intercellular distance of the cell nuclei, play an important role in tissue phenotyping and histologic tumor grading^33,34^. These features can be quantitatively extracted from Deep-CHAMP with high accuracy. Cell counting and nuclear features are derived from FFPE thin tissue slice (Supplementary Fig. 3g,m), fixed and unprocessed thick tissue (Fig. 4i,o), and paraffin-embedded block tissue (Fig. 4v,z). Wilcoxon rank-sum test is applied to evaluate the difference in nuclear features extracted from Deep-CHAMP and gold standard H&E-stained images. The p-value is calculated under a significance level of 0.05. Our results show that the distributions of cross-sectional area and intercellular distance extracted from Deep-CHAMP agree fairly well with the H&E-stained images regardless of the tissue preparation protocols. Although cell counting may be slightly affected by feature segmentation, the distributions of nuclear features still support the accuracy of information that can be extracted from the Deep-CHAMP. These results are highly encouraging and suggest that CHAMP/Deep-CHAMP can be potentially translated into the current histopathological practice to alleviate the workload involved in frozen section or FFPE tissue preparation (Supplementary Fig. 1b).

## Discussion

CHAMP is a promising and transformative histological imaging technology that enables rapid, label-free and high-resolution imaging of thick and unprocessed tissues, holding great promise to streamline the standard-of-care histopathology. However, there are still challenges ahead as CHAMP and H&E-stained images exhibit some deviations. First, the nucleoli structures are better visualized in H&E-stained images than that in CHAMP under the same magnification (e.g., Figs. 2f,k, 4s,w and 5f,h). This is likely because the fluorescence property of nucleoli in the detected spectral range is not chemically identical to H&E histological stains. Second, the densely packed cell nuclei in the hippocampus are less distinguishable in CHAMP compared with the clinical standard images in fresh tissue (e.g., Fig. 5f,h,j,l). This may be attributed to the difference in the imaged thickness (tens of micrometers in CHAMP versus 7 µm in H&E-stained images). Third, fiber tracts in the mouse brain, including corpus callosum and cerebral peduncle, are better visualized in CHAMP than that in H&E-stained images (e.g., Fig. 2b–d,g–i). This is possibly due to the proteins in these fibrous structures present a high quantum yield under deep-UV excitation while eosin exhibits similar affinity as with cytoplasm. Equipping CHAMP with higher optical-sectioning capability is feasible to mitigate the thickness-induced deviations between Deep-CHAMP and H&E-stained images. The axial super-resolution contents can be provided by 3D structured illumination approaches^35,36^, which could potentially facilitate a more comprehensive specimen analysis.

Although an exogenous contrast agent is not required for CHAMP imaging, significant variations in autofluorescence intensity can still be observed in CHAMP as the excitation light is scattered/absorbed differently among various tissue types and functional areas (e.g., white matter and gray matter of the brain)^37^. The fluorescence properties of intrinsic fluorophores, such as excitation/emission maximum and quantum yield, are highly related to the biochemical environment and sample status. A transitory decrease in autofluorescence intensity, which is related to the UV-induced photo-oxidation^38^, can occur at the beginning of UV radiation. With continuous exposure, the intensity is progressively increased, and the resulting homogenously distributed autofluorescence adversely degrades the negative contrast that can be observed in CHAMP images. This is likely because the continuous UV radiation causes damage to the nuclear membrane, thus allowing the efflux of cytoplasmic fluorophores and its binding to nucleolus proteins^28^. Situations are more complicated in fresh tissues, where different levels of local hemorrhage can occur during excision with biopsy forceps, and the consequent attenuation of radiation due to the absorption by non-fluorescent chromophores like hemoglobin^39^ should also be taken into account.

CHAMP can potentially reach higher imaging speed to further shorten the diagnostic timeframe for more time-sensitive applications. Currently, the imaging speed is limited by the exposure time of each speckle-illuminated image, which is nearly 280 ms with an illumination power of 2 mW to maintain a good signal-to-noise ratio (SNR). The acquisition speed can be further accelerated by increasing excitation power. Another limiting factor is the number of acquisitions that are required for super-resolution reconstruction. The minimum number of acquisitions is related to the sparsity of the imaged features. According to our experiments, a sufficiently large scanning range (greater than twice the length of the low-NA diffraction-limited spot size) and fine scanning steps (smaller than the targeted resolution) can reduce distortions in the reconstruction. 2.6× resolution improvement can be obtained through at least 36 speckle-illuminated (average grain size ∼1.3 µm) autofluorescence images without obvious degradation in the reconstructed CHAMP image. The system can be further optimized to balance the achievable resolution and acquisition speed for a specific application. Note that the computational efficiency may be a dominant impediment for CHAMP imaging, which takes ∼50 second/10 mm^2^ due to the use of an iterative reconstruction framework which involves a massive number of Fourier transformations. This issue can be potentially addressed by the implementation of powerful computational resources or the introduction of a deep-learning approach^40^, which allows super-resolution reconstruction with a reduced number of acquisitions under low light conditions. This is expected to dramatically speed up structured illumination approaches and release the computational burden of CHAMP imaging.

It should be admitted that CycleGAN-based network faces difficulty to differentiate features that demonstrate similar brightness (e.g., cell nuclei with interstitial spaces, ventricles, and vessels), which can lead to some staining artifacts (as indicated by the arrows in Supplementary Figs. 3,5). We believe that integrating weakly-/semi-supervised data or introducing a saliency constraint^41^ would help to address this problem, further improving the accuracy of Deep-CHAMP images. Virtual staining through unsupervised learning should be systematically investigated in the future to enable a faithful conversion, which however, is beyond the scope of this study.

In summary, we propose a revolutionary and transformative histological imaging technology which enables rapid, label-free, and high-resolution imaging of thick and unprocessed tissues with large surface irregularity. The versatility of CHAMP is experimentally demonstrated, which enables rapid and accurate pathological examination without tissue processing or staining, demonstrating great potential as an assistive imaging platform for surgeons and pathologists to provide optimal adjuvant treatment intraoperatively. To show diagnostic reliability of CHAMP/Deep-CHAMP, large-scale clinical trials should be carried out as follow-up work. Moreover, computer-aided diagnoses could be incorporated with CHAMP/Deep-CHAMP to further improve the efficiency of the current clinical workflow.

## Methods

### Collection of biological tissues

The organs were extracted from C57BL/6 mice. For fresh tissues (Fig. 5 and Supplementary Fig. 5), the brain/kidney were harvested immediately after the mice was sacrificed and rinsed in phosphate-buffered saline for a few seconds, and then blotted up with laboratory tissue for CHAMP imaging. To prepare fixed and unprocessed tissues (Figs. 3,4a, and Supplementary Fig. 4), the freshly excised tissues were fixed in 4% neutral-buffered formalin at room temperature for 24 hours, and manually sectioned with ∼5-mm thickness or sectioned by a vibratome (VF-700-0Z, Precisionary Instruments Inc.) with ∼200-µm thickness. To prepare paraffin-embedded tissue (Fig. 4p), the formalin-fixed tissues were processed with dehydration, clearing, and infiltration by a tissue processor (Revos, ThermoFisher Scientific Inc.) for 12 hours, and paraffin-embedded as a block specimen. To prepare thin tissue slices (Fig. 2 and Supplementary Fig. 3), the paraffin-embedded block tissues were sectioned at the surface with ∼7-µm thickness by a microtome (RM2235, Leica Microsystems Inc.). The thin tissue slices were stained by H&E, and subsequently imaged by a digital slide scanner (NanoZoomer-SQ, Hamamatsu Photonics K.K.) to generate the histological images. All experiments were carried out in conformity with a laboratory animal protocol approved by the Health, Safety and Environment Office (HSEO) of Hong Kong University of Science and Technology (HKUST).

### Reflection-mode CHAMP system

As shown in Fig. 1a, a nanosecond UV pulsed laser is used as the excitation source (266 nm wavelength, WEDGE HF 266 nm, Bright Solutions Srl.), which is spectrally filtered by a bandpass filter (FF01-300/SP-25, Semrock Inc.) and expanded by a pair of lenses (LA4647-UV and LA4874-UV, Thorlabs Inc.). After that, the expanded beam is obliquely reflected by a UV mirror (PF10-03-F01, Thorlabs Inc.) and projected onto a diffuser (DGUV10-600, Thorlabs Inc.) to generate a constant speckle pattern, which is subsequently focused onto the bottom surface of a specimen by a condenser lens (LA4148-UV, Thorlabs Inc.) with an illumination power of 2 mW. The excited autofluorescence signal is detected by an inverted microscopy system which consists of a plan achromat infinity-corrected objective lens (RMS4X, NA = 0.1, Thorlabs Inc.) and an infinity-corrected tube lens (TTL180-A, Thorlabs Inc.), and finally imaged by a monochrome scientific complementary metal-oxide-semiconductor (sCMOS) camera (PCO edge 4.2, 2048 × 2048 pixels, 6.5-μm pixel pitch, PCO Inc.). In our experiment, the specimen was 2D raster-scanned by a 2-axis motorized stage (L-509.20SD00, PI miCos GmbH) with a scanning interval of 1 µm. A sequence of speckle-illuminated diffraction-limited autofluorescence images were recorded by the sCMOS camera which was synchronized with the motor scanning via our lab-designed LabVIEW software (National Instruments Corp.) and triggering circuits. The overall data acquisition time for each raw image was estimated to be 280 ms (250-ms camera integration time plus 30-ms stage settling time), which was selected to provide a good balance between the acquisition speed and image SNR.

### Super-resolution reconstruction framework

Extended DOF enabled by the implementation of a low-NA imaging objective in the CHAMP microscope not only matches the optical-sectioning thickness provided by UV surface excitation, but also accommodates different levels of tissue surface irregularities. To bypass the resolution limit set by the low-NA objective and maximize the achievable imaging throughput, structured illumination with a constant speckle pattern is implemented. Undetectable high-frequency information can be multiplexed into the low-NA imaging system through intensity modulation, where the fluorescent specimen is illuminated with non-uniform intensity-varied patterns, including sinusoidal stripe^42,43^, multifocal spot^44,45^, or random speckle patterns^46,47–51^. The processes of intensity modulation with uniform illumination, linear/nonlinear sinusoidal stripe, and random speckle pattern are respectively demonstrated in Supplementary Fig. 6. The highest achievable resolution through structured illumination is determined by the reciprocal of Fourier space bandwidth, which is given by *λ*/2(NA_*obj*_ + NA_*illu*_), where *λ* is the fluorescence emission wavelength, and NA_*obj*_ and NA_*illu*_ are the numerical apertures of the detection objective lens and illumination pattern, respectively. The resolution improvement through conventional SIM with epi-fluorescence configuration is restricted to 2× as the illumination NA is also restricted by the detection objective (Supplementary Fig. 6e–h). The adoption of high-frequency sinusoidal harmonics generated by nonlinear fluorescence response allows reaching beyond 2× resolution enhancement^52,53^ (Supplementary Fig. 6i–l). However, the photodamage and photobleaching associated with high-power excitation will hinder its biomedical applications. Recent studies show that 4× resolution improvement can be achieved through an off-axis projection of a set of frequency-multiplexed sinusoidal patterns^54^. However, the system complexity is inevitability increased, and the transmission-based configuration restricts its application only to thin samples.

**Figure 6.**
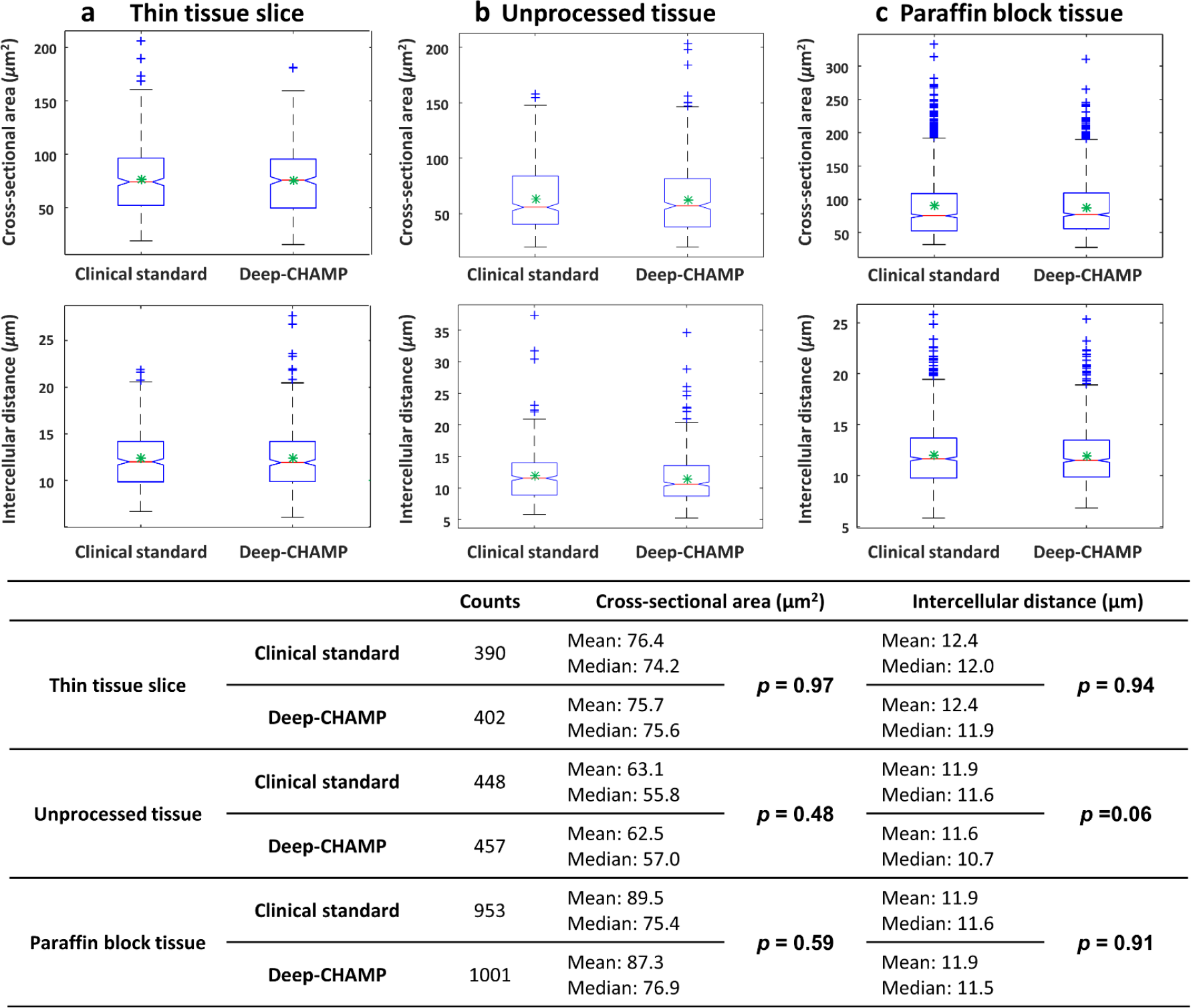
Distributions of nuclear features extracted from Deep-CHAMP and clinical standard images. **a**–**c**, The cross-sectional area and intercellular distance extracted from FFPE thin tissue slice, fixed and unprocessed thick tissue, and paraffin-embedded block tissue, respectively. The table below shows the corresponding cell counts, cross-sectional area, and intercellular distance with statistical analysis. The null hypothesis is accepted as *p* > 0.05.

For simplicity and flexibility, oblique illumination with a constant speckle pattern, which features a grain size smaller than the PSF of the detection optics, is implemented in CHAMP to obtain beyond 2× resolution enhancement. In addition, with the assistance of UV surface excitation, our method is applicable to unprocessed and unlabeled tissues with any thickness, which is not achievable with the existing super-resolution SIM systems. Unlike SIM with sinusoidal illumination, where super-resolution demodulation can be achieved through a one-step analytic inversion with a few raw images, SIM with speckle illumination (e.g., CHAMP, Supplementary Fig. 6m–p) fails to establish a direct inversion relationship, thus requiring more redundant acquisitions to isotropically fill the Fourier space. Note that even with prior information based on speckle statistics^49,55,56^ and sample sparsity^50,57^, multifold resolution gain is not experimentally achievable with fully-randomized speckle patterns due to the ill-posed nature in this situation, i.e., *N* intensity measurements are captured with *N*+1 unknown variables to be solved (*N* illumination patterns plus one sample distribution). To address this issue, illuminating with a constant speckle pattern that is translated between measurements, as opposed to randomly changing speckle patterns, is utilized in this report^46,51,58^.

The CHAMP reconstruction framework is based on a momentum-assisted regularized ptychographic iterative engine^59^, which is a well-developed inversion solver with the significantly improved robustness that enables rapid convergence to a lower error (10 iterations are generally sufficient in our experiments). The flowchart of the reconstruction algorithm is shown in Supplementary Table 1. Before reconstruction, the raw images were flattened to correct illuminance nonuniformity. Then, the scanning trajectory (*x*_*j*_, *y*_*j*_) of the specimen was pre-estimated by cross-correlation of the captured raw images^60^. Note that the sampling rate is a prerequisite for digital image reconstruction. Undersampling issue, which occurs in the CHAMP system as the sampling pixel size is larger than half of the PSF size of the low-NA detection optics, will lead to pixel aliasing and consequently generate artifacts in the reconstruction. To tackle this issue, a sub-sampled method^61^ was introduced. The algorithm was run on a workstation with a Core i9-10980XE CPU @ 4.8GHz and 8×32GB RAM, and 4 NVIDIA GEFORCE RTX 3090 GPUs, which takes ∼50 second/10 mm^2^ for computation.

Note that the resolution improvement is theoretically infinite, which, however, will be experimentally restricted by the speckle contrast on the specimen. In principle, condenser lens with higher illumination NA enables higher achievable resolution at the expense of more acquisitions. However, the resulting highly compressed speckle pattern not only causes vignetting effect which darkens the corners of the captured autofluorescence images, but also degrades the speckle contrast due to the natural decay governed by the incoherent optical transfer function. Therefore, the system can be optimized to balance the tradeoffs between target resolution, acquisition speed and computational efficiency for various applications. In this work, 2.6× resolution gain was achieved via 36 speckle-illuminated (average grain size ∼1.3 µm) diffraction-limited images that were raster-scanned with 1-µm scanning interval. High throughput of 200 megapixels and a long DOF of 80 µm were obtained in CHAMP, enabling rapid and label-free imaging of thick and unprocessed tissues with large irregular surface at an acquisition speed of 10 mm^2^/10 seconds with 1.1-µm lateral resolution (Supplementary Fig. 7).

The spatial resolution of CHAMP was measured by imaging 500-nm-diameter fluorescent beads (B500, excitation/emission: 365/445 nm, Thermo Fisher Scientific Inc.) (Supplementary Fig. 7). The Gaussian-fitted data show that the full width at half maximum is 1.1 µm in the reconstructed CHAMP image while 2.9 µm in the diffraction-limited wide-field image, demonstrating 2.6× resolution enhancement through speckle illumination.

### Virtual staining through unsupervised learning

Supplementary Fig. 8 shows the architecture of the generator and discriminator networks. The objective of CycleGAN contains two types of loss functions — adversarial loss^62^ and cycle consistency loss^31^. For adversarial loss, the objective of the discriminator Y (*D*_*Y*_) is calculated as:

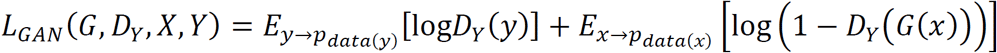

Similarly, the objective of the discriminator X (*D*_*X*_) is:

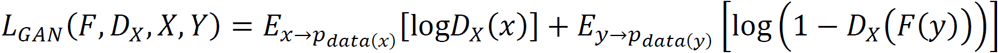

Cycle consistency loss, which is applied to monitor the training process, is calculated as:

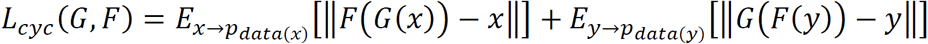

In addition, structural similarity index measure (SSIM)^63^, which predicts the perceived quality based on illuminance, contrast, and structure, is appended to the aforementioned loss functions. The SSIM loss is calculated as:

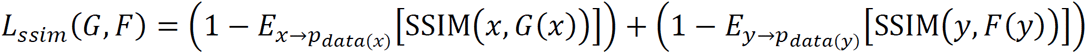

The overall objective for our virtual staining network is the weighted sum of the four loss functions, which is given by:

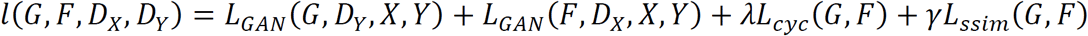

where *λ* is set to 10 and *γ* is set to 2. The network is implemented with Python version 3.7.3 and Pytorch version 1.0.1. The software is implemented on a desktop computer with a Core i7-9700K CPU@ 3.6GHz and 64GB RAM, running on an Ubuntu 18.04.2 LTS operation system. The training and testing are performed by an NVIDIA Titan RTX GPU with 24 GB RAM, which allows operating on ∼25 megapixels/s for testing (including GPU computing and time to write to hard disk).

For virtual staining network of fixed mouse brains, the training data consists of 1,600 unpaired CHAMP and H&E-stained images, where CHAMP images were collectively obtained from fixed, thick/thin mouse brains, and histological images were collected from H&E-stained thin mouse brain slices which contain similar features as the CHAMP images. For virtual staining network of fresh mouse brains, the training data consists of 800 unpaired CHAMP and H&E-stained images, where CHAMP images were collectively obtained from freshly excised mouse brains. For style transformation network, the training data consists of 800 unpaired ‘brain-style’ Deep-CHAMP images and H&E-stained images of thin mouse kidney slices, where Deep-CHAMP images were the output from the fixed mouse brain network with the CHAMP images of mouse kidney used as the input. The training details and convergence plots can be found in Supplementary Fig. 9.

To show the wide applicability of the unsupervised network, the fixed mouse brain/kidney tissues with various thicknesses were utilized for cross validation. The resulting virtually stained Deep-CHAMP images are enumerated in Supplementary Fig. 10. We emphasize that the CycleGAN-based network enables image translation without paired training data, thus fundamentally favouring the virtual staining of CHAMP images of thick and unprocessed tissues. The network was trained with the hybrid CHAMP images of unprocessed/processed, thick/thin tissues that can exhibit differences in terms of cellular morphology, image contrast, and brightness, thus demonstrating strong applicability in different tissue thicknesses (Supplementary Fig. 10i,j). In addition, the bridge network enables style transformation from ‘brain-style’ Deep-CHAMP images (Supplementary Fig. 10k,l) to ‘kidney-style’ Deep-CHAMP images (Supplementary Fig. 10m,n) without the need for retraining a kidney network. Because of this, cross-organ validation with CycleGAN is feasible. This transformation is also applicable to other different types of tissues as long as the CHAMP images of these tissues do not show significant difference with the mouse brain, showing the great simplicity and flexibility of the unsupervised neural network.

### Calculations of cross-sectional area and intercellular distance

Deep-CHAMP and H&E-stained histological images were segmented by a free Fiji plugin, trainable Weka segmentation^64^, which enables to produce pixel-based segmentations. Based on the resulting probability maps, images were subsequently converted to a binary image where cell nuclei can be identified. The binarized Deep-CHAMP and H&E-stained images were analyzed in Fiji, where the cross-sectional area and centroid of each nucleus can be provided. With the localized center positions of the cell nuclei, the intercellular distance was calculated to be the shortest adjacent distance to a neighboring cell nucleus.

## Supporting information

none

## Data availability

All data involved in this work, including raw/processed images provided in the manuscript and supplementary information, are available from the corresponding author upon request.

## Code availability

The customized code in MATLAB for CHAMP super-resolution reconstruction is available at https://github.com/TABLAB-HKUST/CHAMP.

The code of virtual staining networks based on the architecture of CycleGAN is available at https://github.com/TABLAB-HKUST/Deep_CHAMP.

## Acknowledgments

The Translational and Advanced Bioimaging Laboratory (TAB-Lab) at HKUST acknowledges the support of the Hong Kong Innovation and Technology Commission (ITS/036/19); Research Grants Council of the Hong Kong Special Administrative Region (26203619); The Hong Kong University of Science and Technology startup grant (R9421). The authors also appreciate Dr. Siyuan Dong from Massachusetts Institute of Technology and Dr. Li-Hao Yeh from the University of California, Berkeley for the valuable discussion on the reconstruction algorithm.

## Author contributions

Y.Z., L.K., X.F.L. and T.T.W.W. conceived of the study. Y.Z. built the imaging system. L.K. wrote the control software. Y.Z. and L.K. prepared the specimens involved in this study. Y.Z. and I.H.M.W performed imaging experiments. L.K. performed histological staining. Y.Z. processed and analyzed the data. Y.Z. and T.T.W.W. wrote the manuscript. T.T.W.W. supervised the whole study.

