## Supplementary material for "High-throughput, label-free and slide-free histological imaging by computational microscopy and unsupervised learning": none

The Supplementary Material includes:

Supplementary Figure 1 | Workflow of surgical margin assessment.

Supplementary Figure 2 | Comparison of state-of-the-art slide-free imaging modalities.

Supplementary Figure 3 | CHAMP and Deep-CHAMP validation with a thin mouse brain slice.

Supplementary Figure 4 | CHAMP and Deep-CHAMP validation with fixed and unprocessed mouse brain/kidney tissues.

Supplementary Figure 5 | CHAMP and Deep-CHAMP validation with a freshly excised mouse kidney tissue.

Supplementary Figure 6 | Illustration of intensity modulation by structured illumination microscopy.

Supplementary Figure 7 | Experimental characterization of CHAMP's lateral resolution.

Supplementary Figure 8 | Architecture of generator and discriminator neural networks.

Supplementary Figure 9 | Convergence plots and training details.

Supplementary Figure 10 | Cross-validation of the virtual staining network.

Supplementary Table 1 | Flowchart of super-resolution reconstruction framework.

Other Supplementary Material for this manuscript includes the following:

Supplementary Video 1 | System setup, image acquisition, and data processing.

Supplementary Video 2 | A series of close-up and registered CHAMP, Deep-CHAMP, and H&E-stained histological images of a thin mouse brain slice.

Supplementary Video 3 | A series of close-up and registered CHAMP, Deep-CHAMP, and H&E-stained histological images of formalin-fixed thick mouse brain tissues.

Supplementary Video 4 | A series of close-up and registered CHAMP and Deep-CHAMP images of vibratome-cut mouse brain/kidney tissues.

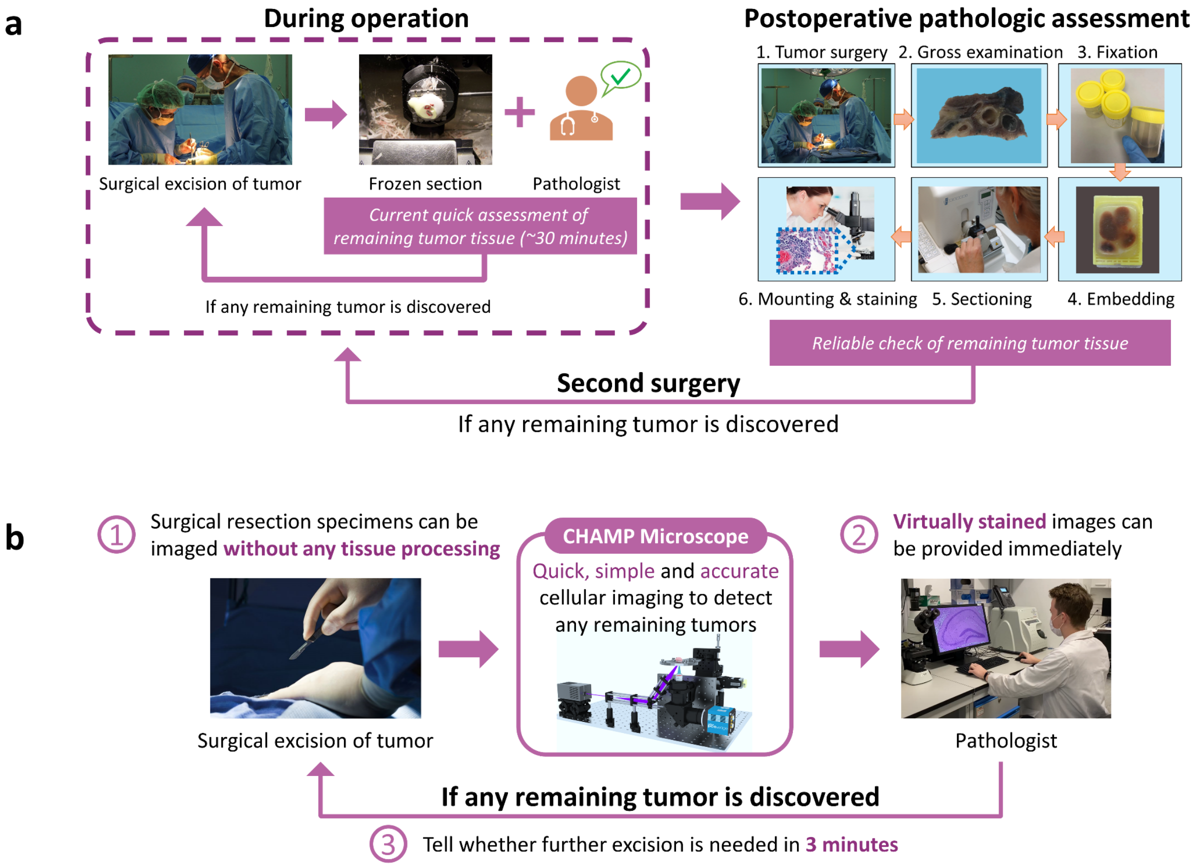


**Supplementary Figure 1 | Workflow of surgical margin assessment. a**, Conventional standard-of-care histopathology, which involves two approaches, including (1) intraoperative frozen section with freshly excised tissues, and (2) postoperative assessment with formalin-fixed and paraffin-embedded tissues. **b**, Anticipated new clinical practice by using the CHAMP microscope.

| Method | Image acquisition | Contrast mechanism | Optical sectioning ability | Imaging throughput | Ease of use | Cost-effectiveness |
|---|---|---|---|---|---|---|
| LSCM | 2D/3D laser scanning | Exogenous/endogenous fluorescence | ★★★★ Controlled by pinhole size, can be traded with SNR and signal intensity | ★★ ~10 megapixels Determined by laser scanning speed and dwell time | ★★ High system complexity in terms of optical alignment, system synchronization, and maintenance | ★★ Require high-speed laser scanning module |
| MPM | | | ★★★★★ Controlled by the size of tightly-focused beam | ★ ~2 megapixels @ 80 MHz laser Determined by laser repetition rate and dwell time | | ★ Require ultrafast pulsed laser with high peak power, and high-speed laser scanning module |
| SRS | | Intrinsic vibration of lipids, proteins, and nucleic acids | | | | |
| SHG | | Non-centrosymmetric properties of endogenously orientated structure | | | | |
| PAM | | Optical absorption of endogenous biomolecules | ★★ Controlled by the bandwidth of ultrasonic transducer | ★★ ~10 megapixels @ 100 kHz laser Determined by laser repetition rate and dwell time | ★★★ Require coupling media and no image can be provided in real time | ★★ Require pulsed laser with high energy, and high-speed laser scanning module |
| LSFM | Wide-field illumination | Two exogenous fluorescence analog of H&E | ★★★★ Controlled by the thickness of light-sheet, can be traded with field-of-view | ★★★★★ ~850 megapixels Determined by exposure time and light sheet thickness | ★★★ Tissue clearing for volumetric imaging; ★★★ Require well-regulated staining protocol to avoid fluorescence saturation and leakage | ★★★ Require laser and high-speed camera |
| SIM | | | ★★★ Controlled by the frequency of illumination pattern, can be traded with SNR and imaging depth | ★★★★★ ~800 megapixels Determined by exposure time and pattern switching time | ★★★ Precisely controlled pattern generation module | ★★★★ Require spatial light modulator and camera with high dynamic range |
| MUSE | | | ★★ Controlled by the UV penetration depth, which is tissue dependent | ★★★★ ~110 megapixels Determined by exposure time and number of axial scanning for extended DOF | ★★★★ Variable focusing is required for surface irregularities | ★★★★★ Only require UV LED |
| CHAMP | | Endogenous fluorescence | | ★★★★ ~200 megapixels Determined by exposure time and number of acquisitions required for super-resolution reconstruction | ★★★★★ Label-free, large DOF accommodates tissue irregularities | ★★★★ Require UV laser |

**Supplementary Figure 2 | Comparison of state-of-the-art slide-free imaging modalities.** The throughput of each imaging modality is calculated based on the reported literature, including laser scanning confocal microscopy (LSCM)[1], multiphoton microscopy (MPM)[2], stimulated Raman scattering (SRS) and second harmonic generation (SHG)[3], photoacoustic microscopy (PAM)[4], light-sheet fluorescence microscopy (LSFM)[5], structured illumination microscopy (SIM)[6], and microscopy with ultraviolet surface excitation (MUSE)[7].

* Note. Throughput is defined by the ratio of attainable field-of-view per minute to the square of half-pitch resolution. For an easy comparsion, computational time and fluorescence labeling time are not considered here. Throughput is calculated for each imaging modality under the same level of tissue irregularity of 80 µm. For instance, in LSCM[1], 10 $mm^2$/minute with 0.6-µm lateral resolution and 8-µm focus tracking is achieved, such that the throughput is calculated as $\frac{10\ \mathrm{mm}^2/(0.6\ \mu\mathrm{m}/2)^2}{(80\ \mu\mathrm{m}/8\ \mu\mathrm{m})} \approx 10$ megapixels. In nonlinear microscopy, including MPM[2], SRS, and SHG[3], 1 $mm^2$/minute with 0.4-µm lateral resolution is obtained, and the axial scanning interval is 5 µm for each slice, such that the throughput is calculated as $\frac{1\ \mathrm{mm}^2/(0.4\ \mu\mathrm{m}/2)^2}{(80\ \mu\mathrm{m}/5\ \mu\mathrm{m})} \approx 2$ megapixels. In LSFM[5], 100 $mm^2$/12.5 s at 80-µm depth-of-field (DOF) with 1.5-µm lateral resolution is achieved, such that the throughput is calculated as $480\ \mathrm{mm}^2/(1.5\ \mu\mathrm{m}/2)^2 \approx 850$ megapixels. In MUSE[7], 110 $mm^2$/minute with 0.7-µm lateral resolution is achieved, and multiple z-stacks are acquired at 10-µm spacing for extended DOF, such that the throughput is calculated as $\frac{110\ \mathrm{mm}^2/(0.7\ \mu\mathrm{m}/2)^2}{(80\ \mu\mathrm{m}/10\ \mu\mathrm{m})} \approx 110$ megapixels. In CHAMP, 60 $mm^2$/minute with 1.1-µm lateral resolution at 80-µm DOF is achieved, such that the throughput is calculated as $60\ \mathrm{mm}^2/(1.1\ \mu\mathrm{m}/2)^2 \approx 200$ megapixels.

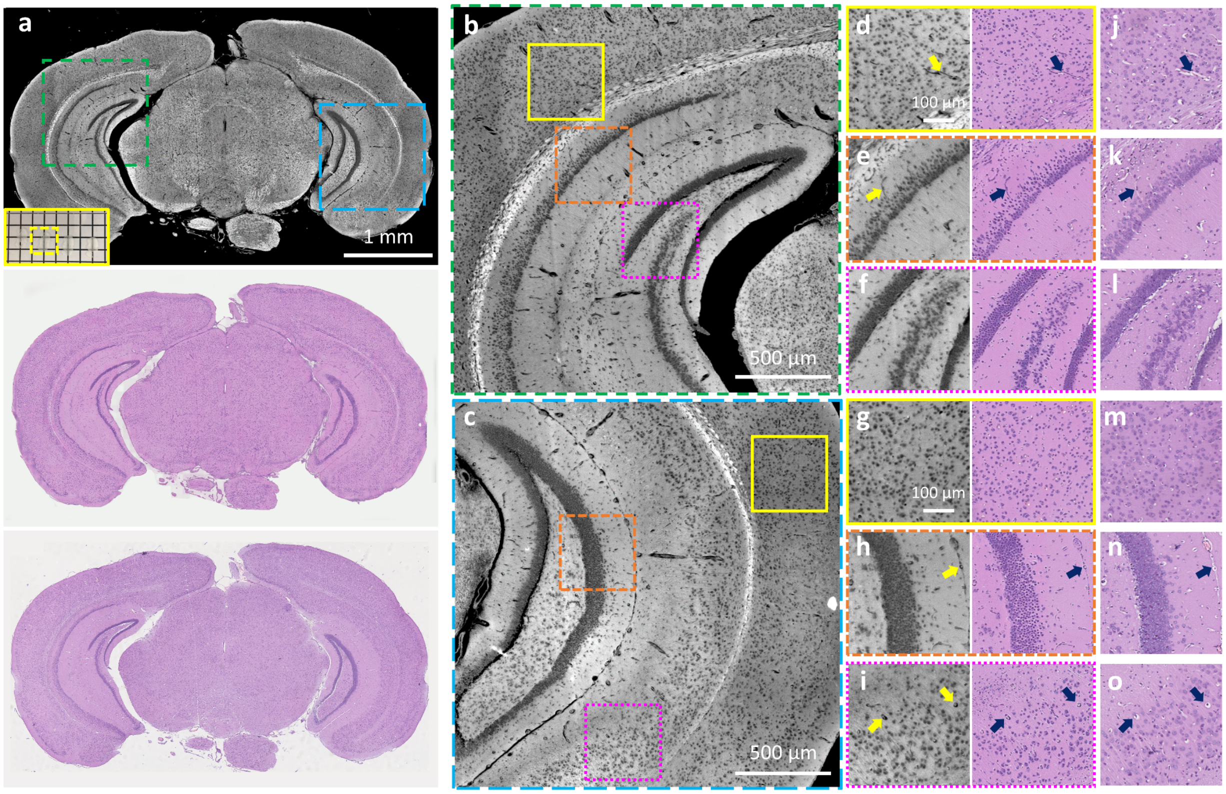


**Supplementary Figure 3 | CHAMP and Deep-CHAMP validation with a thin mouse brain slice. a**, CHAMP (top), Deep-CHAMP (middle), and H&E-stained image (bottom) of a thin mouse brain slice, inset at the bottom left of CHAMP shows the photograph of the specimen (the yellow dashed box shows the mouse brain slice that is imaged). **b**,**c**, Zoomed-in CHAMP images of green and blue dashed regions in a, respectively. **d**–**f**,**g**–**i**, Zoomed-in CHAMP and Deep-CHAMP images of yellow solid, orange dashed, and magenta dashed regions in b and c, respectively. **j**–**o**, The corresponding H&E-stained histological images. Arrows indicate the segmentation-induced staining artifacts in Deep-CHAMP images.

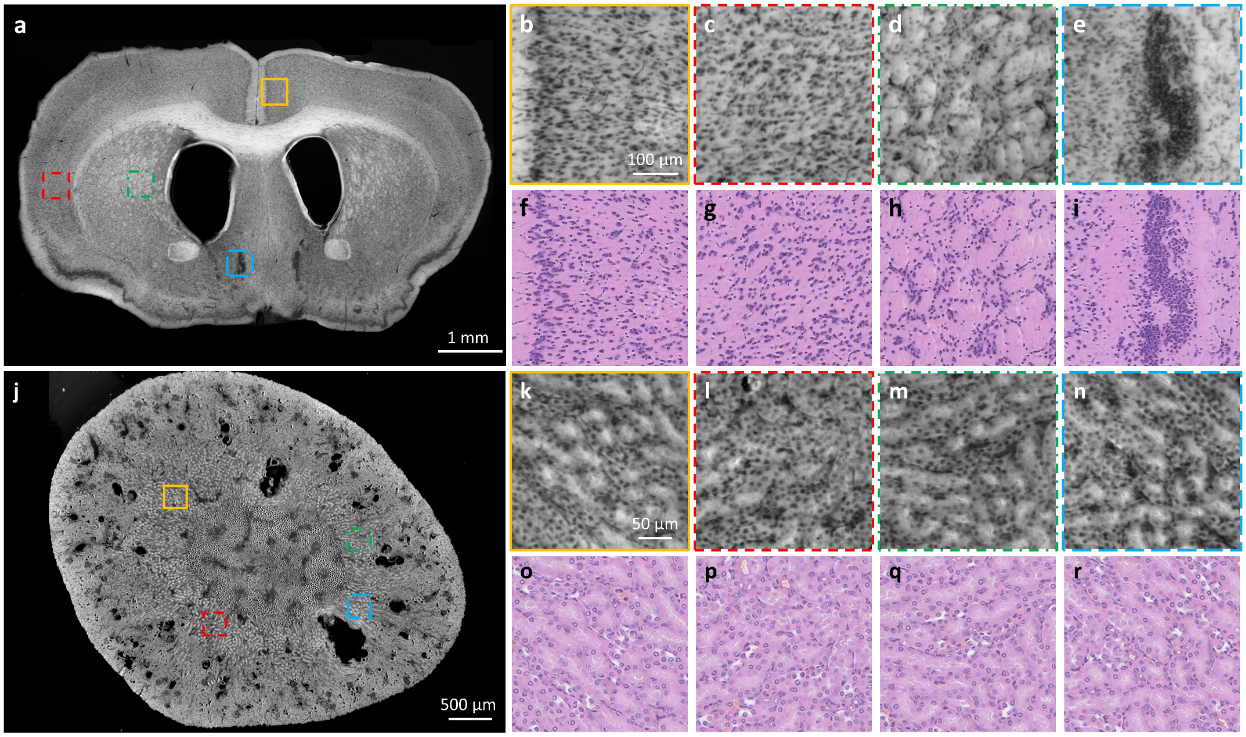


**Supplementary Figure 4 | CHAMP and Deep-CHAMP validation with fixed and unprocessed mouse brain/kidney tissues. a**, CHAMP image of a fixed and unprocessed mouse brain with thickness ~200 µm. **b**–**e**, Zoomed-in CHAMP images of orange solid, red dashed, green dashed, and blue dashed regions in a, repectively. **f**–**i**, The corresponding virtually stained Deep-CHAMP images. **j**, CHAMP image of a fixed and unprocessed mouse kidney with thickness ~200 µm. **k**–**n**, Zoomed-in CHAMP images of orange solid, red dashed, green dashed, and blue dashed regions in j, respectively. **o**–**r**, The corresponding virtually stained Deep-CHAMP images.

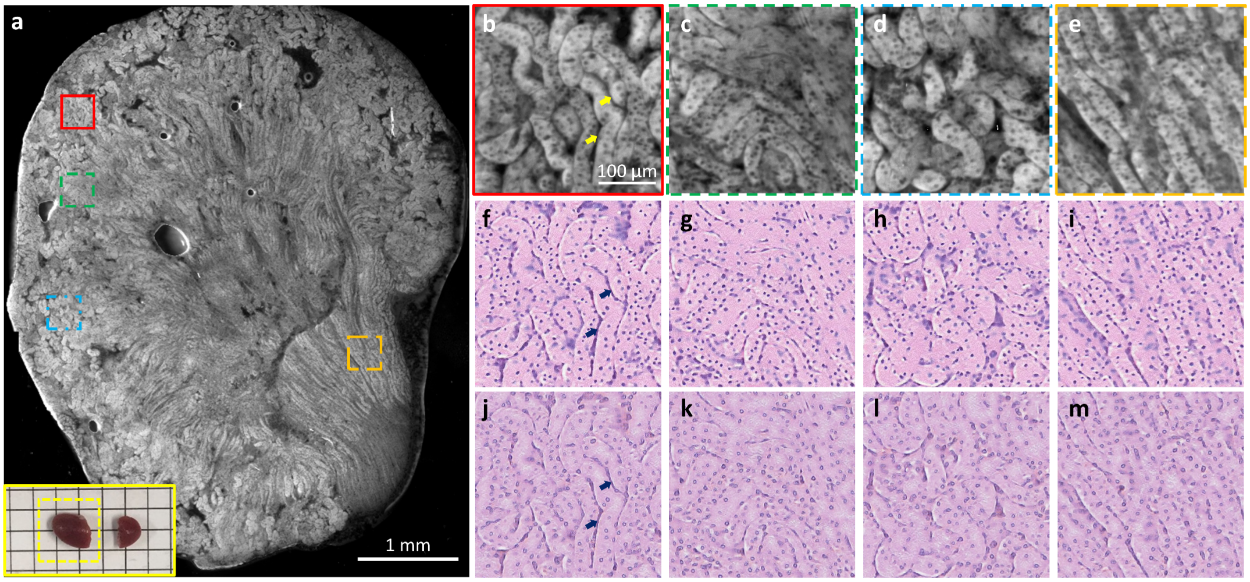


**Supplementary Figure 5 | CHAMP and Deep-CHAMP validation with a freshly excised mouse kidney tissue. a**, CHAMP image of a freshly excised mouse kidney tissue, inset at the bottom left shows the photograph of the specimen (the yellow dashed box shows the mouse kidney that is imaged). **b**–**e**, Zoomed-in CHAMP images of red solid, green dashed, blue dashed, and orange dashed regions in a, respectively. **f**–**i**, The corresponding 'brain-style' Deep-CHAMP images output by the virtual staining network trained for fresh mouse brain. **j**–**m**, The corresponding 'kidney-style' Deep-CHAMP images output by the style transformation network with f–i as the input. Arrows indicate the segmentation-induced staining artifacts in Deep-CHAMP images.

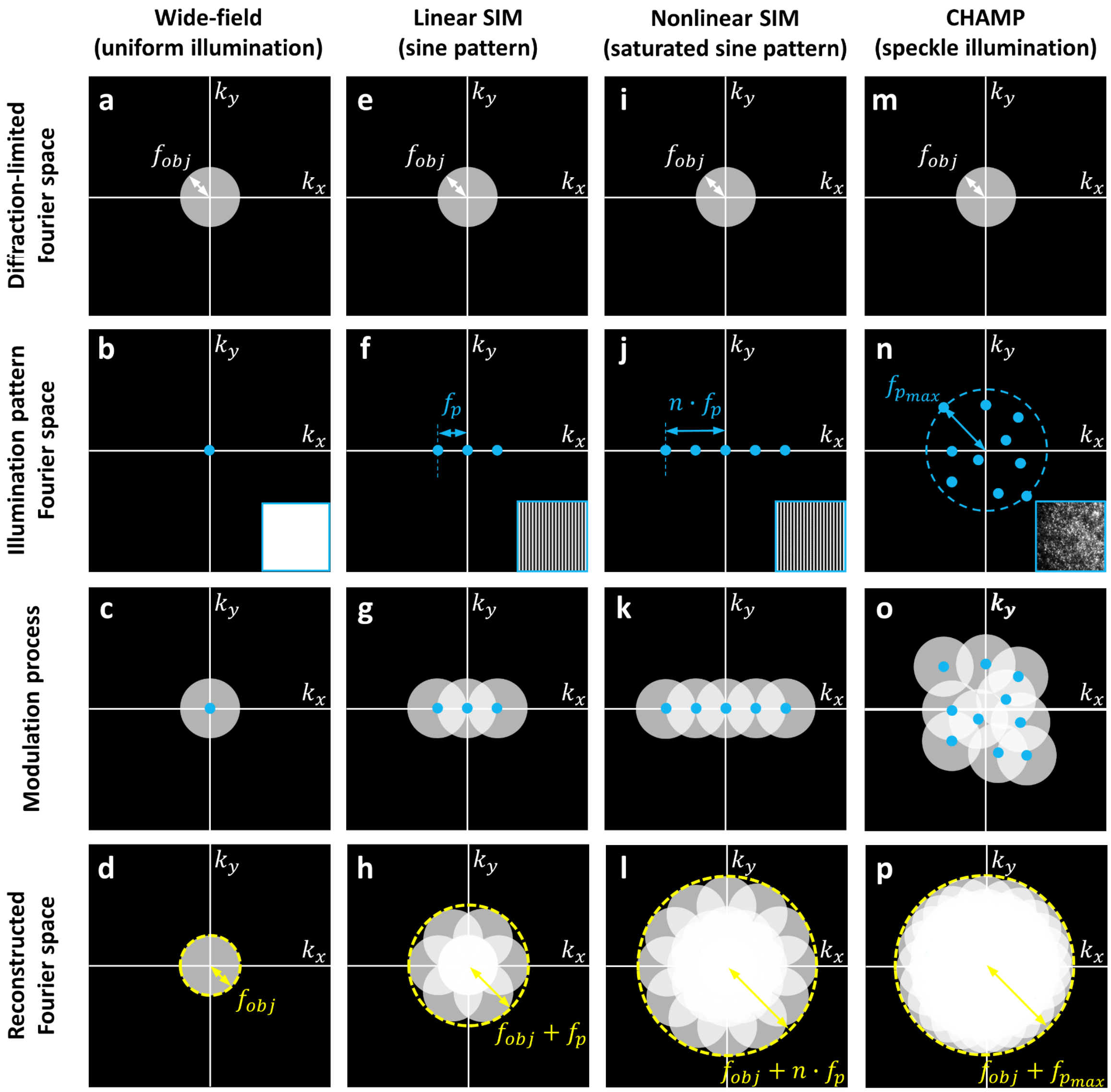


**Supplementary Figure 6 | Illustration of intensity modulation by structured illumination microscopy.** **a**–**d**, Diffraction-limited wide-field imaging with uniform illumination. **e**–**h**, Linear SIM with sinusoidal illumination. **i**–**l**, Nonlinear SIM with saturated sinusoidal illumination. **m**–**p**, CHAMP imaging with translated speckle illumination. For SIM, the sinusoidal pattern is phase-shifted and rotated to synthesize an isotropic aperture (h,l). While for CHAMP, the speckle pattern is translated to isotropically fill the Fourier space (p). $f_{obj}$: frequency of objective lens, $f_p$: frequency of sinusoidal pattern, $n$: the order of sinusoidal harmonics, $f_{p_{max}}$: the maximum frequency of the speckle pattern.

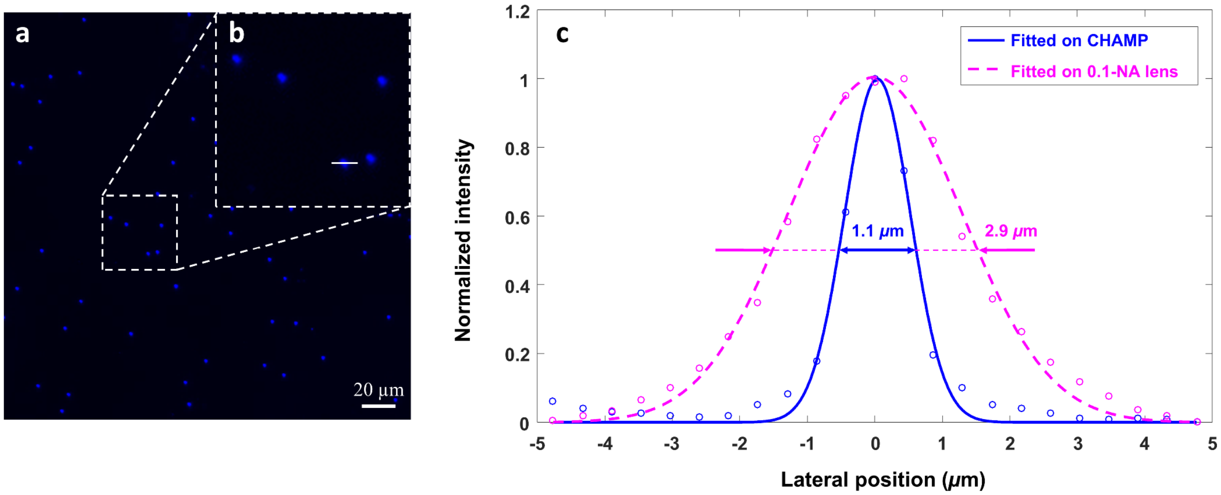


**Supplementary Figure 7 | Experimental characterization of CHAMP's lateral resolution. a**, CHAMP image of blue fluorescent beads (500-nm in diameter with an emission wavelength of 445 nm). **b**, Zoomed-in CHAMP image of the white dashed box in a. **c**, Gaussian-fitted intensity distribution along the solid line in b, showing that the full width at half maximum is 1.1 µm in CHAMP (blue solid line) and 2.9 µm in wide-field microscopy with a 0.1-NA objective lens (magenta dashed line).

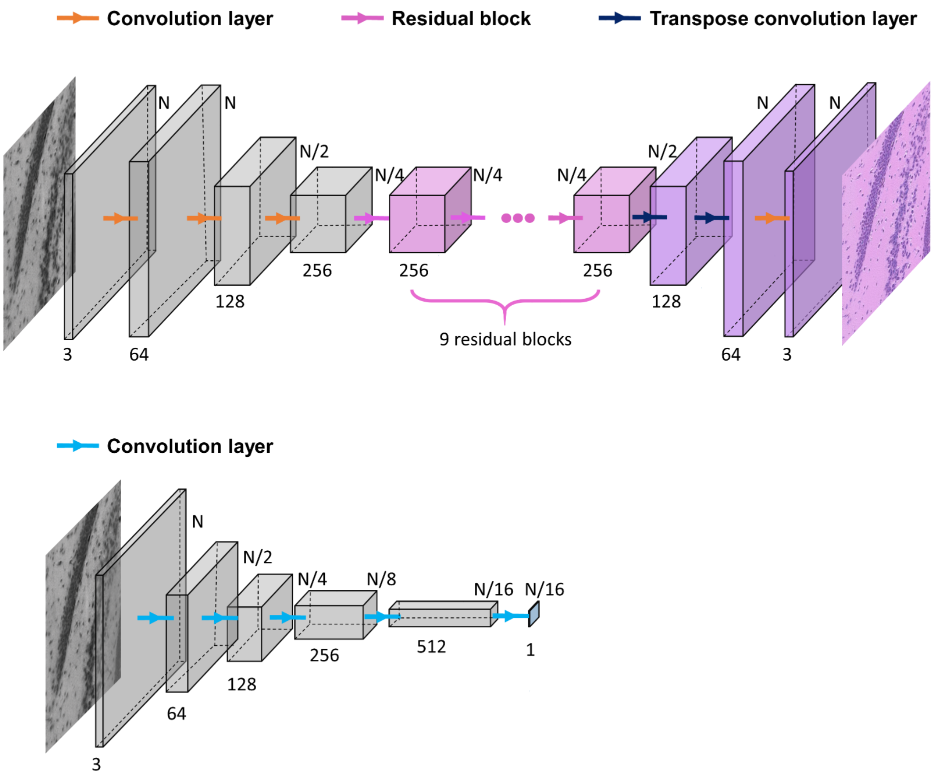


**Supplementary Figure 8 | Architecture of generator and discriminator neural networks.** The Resnet-based generator network[8] (top) consists of a downsampling path (gray), a residual path (pink), and an upsampling path (purple). The first convolution layer (kernel 7 × 7, stride 1 × 1) in the downsampling path increases the image channel to 64 with size unchanged, while the other two layers (kernel 3 × 3, stride 2 × 2) will halve the image size and double the image channel. The network is then followed by 9 residual blocks, in which the image size and channel will remain unchanged. In the upsampling path, the first two layers (kernel 3 × 3, stride 2 × 2) will double the image size and halve the image channel, while the third layer (kernel 7 × 7, stride 1 × 1) decreases the image channel to 3. The last convolution layer is followed by a hyperbolic tangent activation while other convolution layers are followed by an instance normalization and rectified linear unit (ReLU) activation. The PatchGAN-based discriminator network[9] (below) consists of 5 convolutional layers. The image size will be halved by each of the first 4 convolution layers (kernel 4 × 4, stride 2 × 2), with each layer followed by an instance normalization and Leaky-ReLU activation. The last layer (kernel 4 × 4, stride 1 × 1) decreases the channel to 1 to output the probability labels.

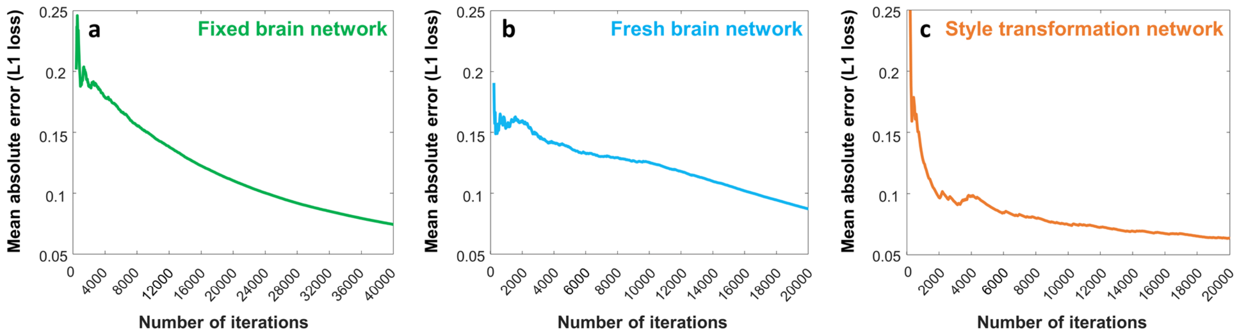


| Network | # of epochs | # of iterations | # of patches (256 × 256 pixels) | Batch size | Training time (h) |
|---|---|---|---|---|---|
| Fixed brain network | 100 | 40000 | 1600 | 4 | 8 |
| Fresh brain network | 100 | 20000 | 800 | 4 | 4 |
| Style transformation network | 100 | 20000 | 800 | 4 | 4 |

**Supplementary Figure 9 | Convergence plots and training details. a–c**, L1-loss (moving average) with respect to the number of iterations of the fixed brain network, fresh brain network, and style transformation network, respectively. The table below shows the parameters of the neural networks a–c.

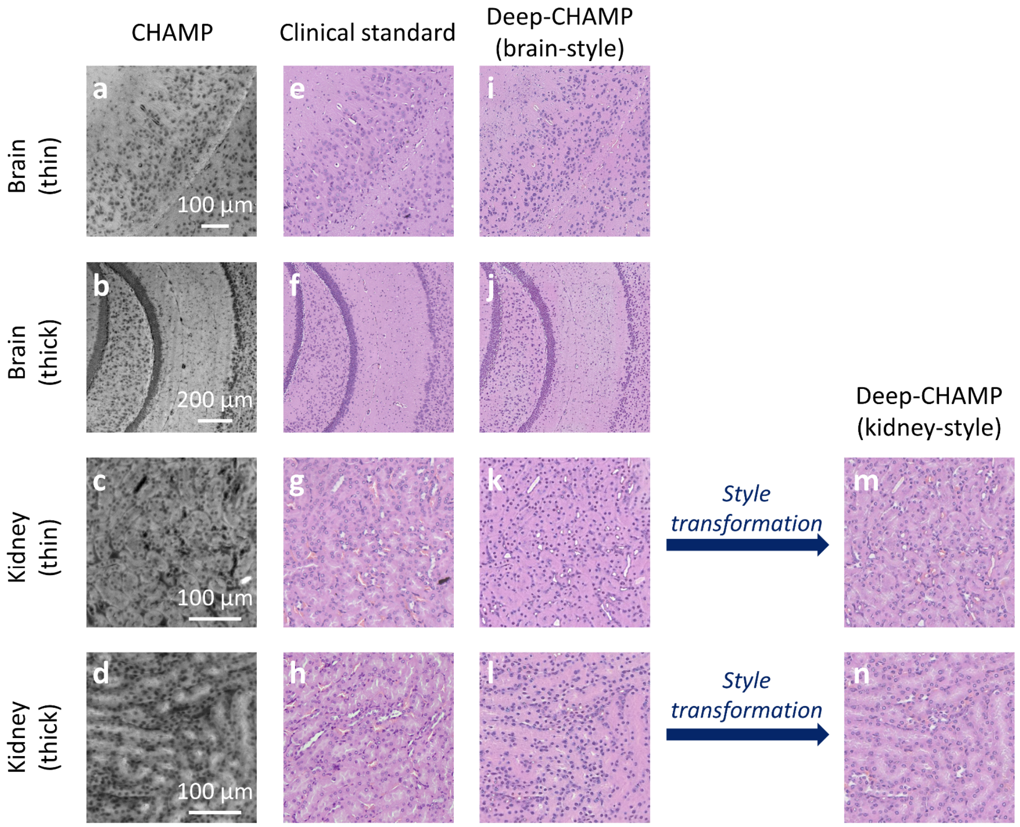


**Supplementary Figure 10 | Cross-validation of the vitual staining network. a–d**, CHAMP images of formalin-fixed mouse brain/kidney tissues with varying thickness. **e–h**, The corresponding H&E-stained images. **i–l**, The corresponding 'brain-style' Deep-CHAMP images output by the virtual staining network for fixed mouse brains. **m**,**n**, The corresponding 'kidney-style' Deep-CHAMP images output by the style transformation network with k and l as the input, respectively.

**Reconstruction Framework**

Input: 36 speckle-illuminated autofluorescence images $I_j$ and pre-estimated position shifts $(x_j, y_j)$
Output: Resolution-enhanced object $o(x, y)$ and unknown speckle pattern $p(x, y)$

**1.** Initialize $o(x, y)$ and $p(x, y)$
**2.** for iteration $= 1{:}\,10$
**3.** for $j = 1{:}\,36$
**4.** $o_j = o(x - x_j, y - y_j)$
**5.** $\varphi_j(x, y) = o_j \cdot p(x, y)$
**6.** $\psi_j(k_x, k_y) = F\left(\varphi_j(x, y)\right) \cdot OTF(k_x, k_y)$
**7.** $I_{est} = \left|F^{-1}\left(\psi_j(k_x, k_y)\right)\right|$
**8.** $I_{est}^{update}(1{:}\,3{:}\,3\mathrm{M}, 1{:}\,3{:}\,3\mathrm{M}) = I_j$
**9.** $\mathrm{F}(\varphi_j^{update}) = F(\varphi_j) + \frac{conj(OTF) \cdot \left[F\left(I_{est}^{update}\right) - \psi_j\right]}{(1-\alpha)|OTF|^2 + \alpha|OTF|_{max}^2}$
**10.** $o_j = o_j + \frac{conj(p) \cdot \left(\varphi_j^{update} - \varphi_j\right)}{(1-\beta)|p|^2 + \beta|p|_{max}^2}$
**11.** $p = p + \frac{conj(o_j) \cdot \left(\varphi_j^{update} - \varphi_j\right)}{(1-\gamma)|o_j|^2 + \gamma|o_j|_{max}^2}$
**12.** $o(x, y) = o_j(x + x_j, y + y_j)$
**13.** end
**14.** end

**Supplementary Table 1 | Flowchart of super-resolution reconstruction framework.** The object $o(x, y)$ is firstly initialized by averaging all captured raw images which are correspondingly back-shifted according to the pre-estimated scanning trajectory $(x_j, y_j)$, and followed by zero-padding in the Fourier domain from the size of M × M to 3M × 3M. Similarly, the speckle pattern $p(x, y)$ is initialized by averaging all captured raw images and padded in the Fourier domain to a size of 3M × 3M. For the $j^{th}$ captured image, the pattern $p(x, y)$ is multiplied with a shifted object $o(x - x_j, y - y_j)$ and Fourier transformed into the frequency domain, which is subsequently lowpass filtered by the optical transfer function of the imaging system, and inversely Fourier transformed to obtain an estimated output intensity $I_{est}$. This estimated intensity is updated by the correspondingly captured autofluorescence intensity $I_j$ in the spatial domain with a sub-sampled method to bypass the resolution-limit set by the physical pixel size. After that, the object $o_j$ and speckle pattern $p$ are alternately updated with the momentum-assisted regularized ptychographic iterative engine[10]. We adopt $\alpha = 1$ while $\beta = \gamma = 0.3$ in the updating function. The shifting operation (line 4 and line 12) is achieved by applying the angular spectrum in the frequency domain.
